# Cross-modality Synthesis of EM Time Series and Live Fluorescence Imaging

**DOI:** 10.1101/2022.02.11.480136

**Authors:** Anthony Santella, Irina Kolotuev, Caroline Kizilyaprak, Zhirong Bao

## Abstract

Analyses across imaging modalities allow the integration of complementary spatiotemporal information about brain development, structure and function. However, systematic atlasing across modalities is limited by challenges to effective image alignment. We combine highly spatially resolved electron microscopy (EM) and highly temporally resolved time-lapse fluorescence microscopy (FM) to examine the emergence of a complex nervous system in *C. elegans* embryogenesis. We generate an EM pseudo time series at four classic developmental stages and create a landmark-based co-optimization algorithm for cross-modality image alignment, which handles developmental heterochrony among datasets to achieve accurate single-cell level alignment. Synthesis based on the EM series and time-lapse FM series carrying different cell-specific markers reveals critical dynamic behaviors across scales of identifiable individual cells in the emergence of the primary neuropil, the nerve ring, as well as a major sensory organ, the amphid. Our study paves the way for systematic cross-modality analysis in *C. elegans* and demonstrates a powerful approach that may be applied broadly.

**Highlights:**

- An EM time series to examine the emergence of an entire nervous system
- A landmark-based co-optimization algorithm for cross modality image alignment in the presence of developmental heterochrony
- Integration of EM and fluorescence series reveals cell behaviors at high spatial and temporal resolution
- Systematic single-cell annotation of EM data enables efficient navigation and targeted re-imaging

## Introduction

Imaging modalities, both evolving established methods like Electron Microscopy (EM)^1,2^ and fluorescence microscopy (FM)^3,4^ and emerging methods like spatial sequencing^5^ and micro CT ^6^ provide unprecedented streams of data. This data has revolutionized our ability to observe the development, structure, and function of the nervous system. However, interpretation and synthesis of this data remain challenging. Combining multiple modalities typically provides complementary information yielding a more holistic view of structural and functional dynamics across spatial and temporal scales ^7,8^. Despite this, systematic atlasing that explicitly combines modalities ^9^ is limited, in part due to the technical difficulties in cross-modality image alignment. Most automated alignment methods such as those used in brain and whole organism atlases focus on appearance-based approaches ^9-14^ which align areas with similar appearances. However, in cross-modality applications there is generally little shared appearance. Another class of alignment methods use anatomical landmarks, typically represented as points, and take known correspondences ^15^ or match points ^16-22^ to compute an alignment. To account for complex variation of landmark positions ^18,23^, a variety of approaches are used to exploit relative position to improve matching accuracy ^19-22^, but performance is limited with dense landmarks (typically 50-60% accuracy). Efforts to build in additional information such as multi-color labeling to distinguish the landmarks, while technically difficult, can greatly improve matching performance (to 81-86%) ^17,21^.

Developmental data, which encompass dynamic anatomical changes over an extended time period, contain even more challenges. Different developmental processes show slight variation of developmental speed, creating heterochrony among anatomical structures. Between samples of comparable developmental stage, certain cell types and tissues/organs may or may not be present, causing a mismatch in the landmarks present. These potentially significant differences between samples complicate alignment and matching. More quantitatively, heterochrony in cell movement (from active migration or tissue shape change) may cause systematic positional variation that confounds approaches with naïve assumptions. Alignment using manually selected landmarks that are always present across samples partially bypasses these challenges ^24^, but generally, heterochrony and the associated challenges in developmental data are unaddressed.

Here, we address these limitations in a cross-modal study of *C. elegans* neural development. We build an EM pseudo time series of *C. elegans* embryonic development and conduct cross-modality analysis with time lapse FM to examine neural development with simultaneous high spatial and temporal resolution. EM studies in *C. elegans* have made profound contributions to neuroscience including the first connectome ^25^, sexual dimorphism ^26^, variability among individuals ^27,28^, and principles of structural organization ^26-29^. However, these efforts are focused on the larval and adult nervous system, which leaves unanswered many questions about how the complex structure of the nervous system arises. Since cells are typically identified in EM based on specialized adult morphologies it is challenging to identify cells in the embryo during their emergence. Instead, significant work using live FM has examined *C. elegans* embryogenesis, providing insight into neurulation ^30,31^, organogenesis ^32,33^, neuropil formation ^29,34-37^, synaptic specificity ^38^, as well as lineage differentiation and brain asymmetry ^39-41^. Lineage tracing in FM establishes definitive identities for cells via ancestry even when they lack distinctive positions or morphologies ^35,42^ and light sheet microscopy allows imaging to extend into embryonic motion ^43^. However, these efforts are limited by the lack of spatial resolution and availability of cell-specific markers needed to resolve individual cell shape. To fill in this gap, we generate an EM embryonic series, develop and apply a landmark-based co-optimization algorithm for cross-modality alignment in the presence of developmental variations. Furthermore, we synthesize the EM series and live FM series data representing over a dozen markers. The cross-modality synthesis reveals critical dynamic behaviors of identifiable individual cells across scales in the emergence of the primary neuropil, the nerve ring, as well as the sensory end of a major sensory organ, the amphid. Our study paves the way for systematic cross-modality analysis in *C. elegans* and demonstrates a powerful approach that may be applied broadly. We present the resulting annotated series as a resource for the community in an online format that facilitates additional community-based curation.

## Results

### Multi-modal Developmental Time Series of *C. elegans* Embryogenesis

We first image a dense EM pseudo time series of *C. elegans* embryogenesis with data at four classic stages, the bean, comma, 1.5-fold and 2-fold stages ^44^ (Fig. 1). The EM series spans a critical 150 min time window during which most tissues undergo major morphological changes: pharyngeal and intestinal lumens develop, distinctive morphologies develop in muscles and the excretory system, skin/hypoderm spreads and closes over the embryo, and the body length doubles. In the nervous system, most of the major nerve tracts emerge. The nerve ring is not yet present at the first time point, but densely populated at the end of this period. As part of the effort to test and demonstrate the feasibility of EM time series acquisition, the bean and comma stage embryos are imaged with Focused Ion Beam-Scanning Electron Microscopy (FIB-SEM) while the 1.5 and 2-fold stage embryos are imaged with Array Tomography (AT). The lateral resolution ranges from 8.5 to 19.4 nm, and z spacing 25 to 85 nm (see Table S1 and Methods). Entire embryos were imaged, containing 576-601 cells including visible apoptotic cells (Table S2).

**Figure 1:**
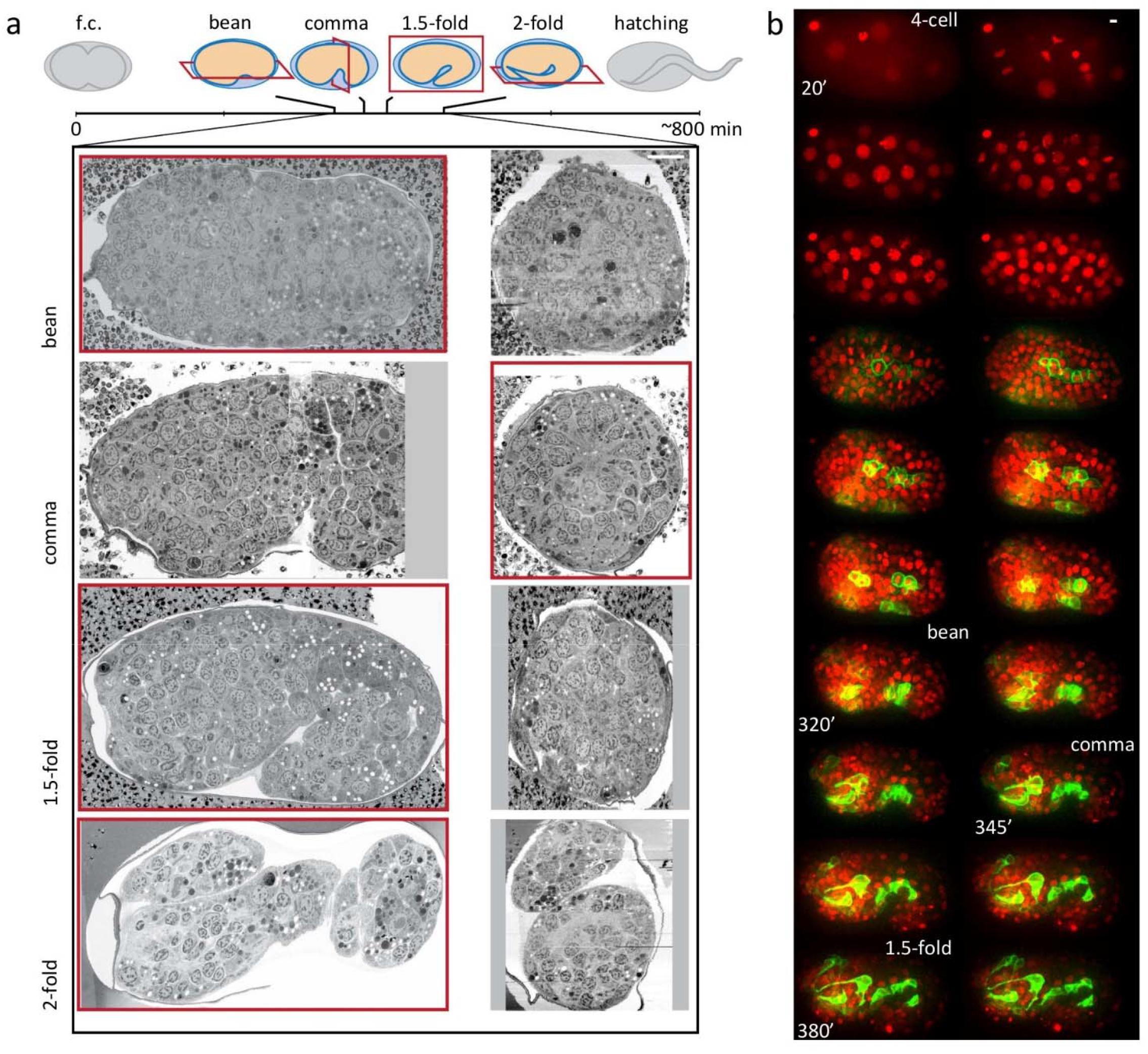
EM and FM Time Series of *C. elegans* Embryogenesis with Complementary Spatial and Temporal Resolution. **(a)** Overview of EM data and their placement in the time course of *C. elegans* embryogenesis. Cartoons show body shape and the approximate physical sectioning plane orientation is shown in red. Orthogonal views of each dataset are shown below, with the view corresponding to physical sectioning outlined in red. f.c., first cell division. **(b)** An example FM series with a ubiquitous histone marker (red) and a *cnd-1* promoter driven membrane marker labeling a subset of neurons (green). Representative time points are shown as max projection. Time stamps are min post first cell division (pfc). Scale bars indicates 5 μm.

3D time-lapse FM is widely used to study *C. elegans* embryogenesis, providing rich information for cross-modality synthesis with EM. We focus on two-color FM images from the WormGUIDES project ^29,45^, where a ubiquitously expressed histone/nuclear marker is used to label and track every cell and a sparsely expressed cell membrane marker reveals dynamic cell behaviors such as neurite outgrowth (Fig. 1b, S1) ^29,46^. These image series typically begin at the 4-cell stage and continue for approximately 8 hours until the 1.75-fold stage and the onset of embryonic movement, with a temporal resolution of 60 or 75 seconds. The nuclear labels are used to trace the entire cell lineage and provide the position and lineage identity of every cell at every time point ^47,48^.

In order to correlate a FM series to the EM series, we use the nuclear positions, which are also annotated for the EM series, as the common reference (i.e., a Rosetta stone). We find the optimal 4D (3D+time) alignment between the FM and EM series based on nuclear positions, which conveys lineage identities to cells in the EM series. We then relate the dynamic cell behaviors observed in the FM and EM series based on the correlated cell identities.

### The Co-optimization Algorithm for Cross-modality Alignment of Developmental Images

We present a landmark-based approach for cross-modality image alignment, which accounts for developmental variation and functions well with challenging dense landmarks. Alignment here takes the form of assigning landmark identities from a set of labeled data to novel unlabeled data, this involves both the computation of a geometric alignment proper and matching. The landmarks, which could be of any anatomical scale that originate from any imaging modality and segmentation method, are represented as spatial point clouds. As detailed below, our algorithm involves an ensemble model of example data to represent the anatomical structure of landmarks in the labeled data and a co-optimization approach for alignment.

The ensemble model contains three parts (Fig. 2a, S4a) in order to capture the regularity and developmental variations of the anatomy: the set of labeled data; a neighbor graph representing consistent adjacencies observed between landmarks across individual labeled datasets; and the set of variable landmarks whose presence is inconsistent among labeled data. First, the ensemble, like instance-based learning approaches that use training data to directly model a distribution, can better preserve the complex positional variation observed in the labeled data. Second, the consistent adjacencies graph summarizes the structural consistency in three dimensional spatial relationships. The adjacency graph captures relative positions, which makes it insensitive to body size variation, or shifting between organs and tissues due to posture, sample distortion or heterochrony in development, since the invariant positional relationships within a given organ/tissue are preserved in the graph. The approach is further motivated by biology. Physical adhesion between cells and tissues underlie anatomical regularity. Consistent adjacencies over multiple samples enrich for the *bona fide* regularity caused by some underlying physical driver and removes coincidental adjacencies. Third, and most importantly, the set of inconsistent landmarks provides a data-driven approach to handle discrepancies in landmarks between labeled and unlabeled data, which has been a major challenge and source of error in alignment. Inconsistent landmarks are likely discrepancies, which can be used to deliberately modify a labeled dataset by adding or removing these landmarks to reduce discrepancies and improve the success of alignment and identity assignment.

**Figure 2:**
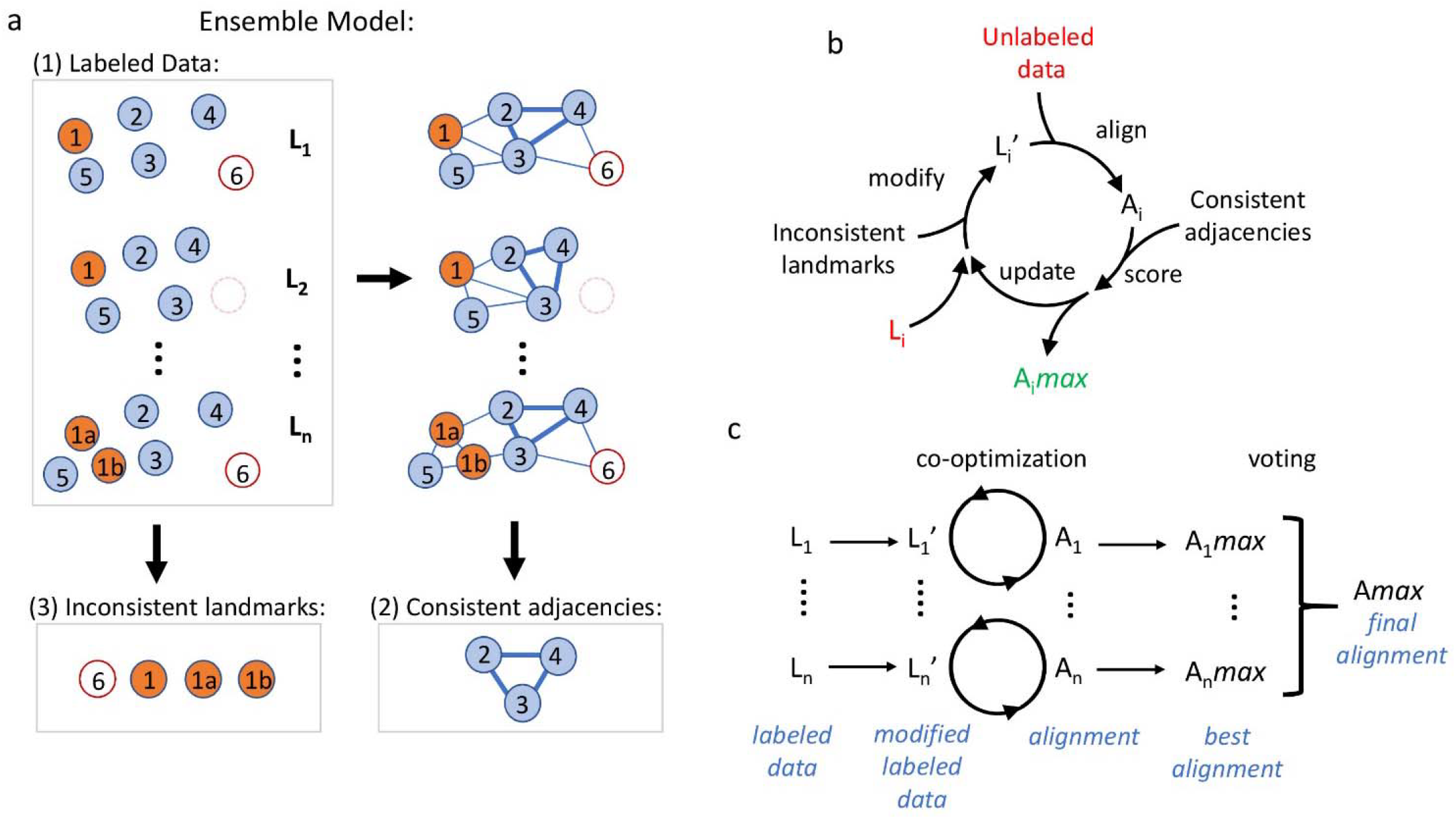
The Co-optimization Algorithm for Cross-modality Alignment of Developmental Data. **(a)** Components of the ensemble model. Circles represent landmarks. Numbers in circle mark the corresponding landmarks across individual label data (L_1_ to L_n_). 1a and 1b denote a split of 1 (cell division or other separation of one landmark into two) and empty circle disappearance of 6 (cell death or other transient landmark disappearance). Lines represent computed adjacencies, among which thick lines represent consistent adjacencies. **(b)** Co-optimization of landmark composition in a labeled dataset L_i_ and alignment A_i_ between L_i_ and unlabeled data. L_i_ is iteratively modified to L_i_’, aligned, and scored to judge each modification. Modifications that improve score are accumulated to yield an optimized L_i_’ and corresponding alignment A_i_max. **(c)** The overall process of deriving correlated identities for an unlabeled dataset. Each labeled dataset L_i_ is optimized to generate a best corresponding A_i_max. Voting per landmark is then performed over the set of labels across A_i_max to achieve a final labeling Amax.

The co-optimization algorithm (Fig. S2) uses the ensemble model to simultaneously optimize modifications to the set of landmarks in labeled data and the alignment between the modified labeled and the unlabeled data (Fig. 2b). Adding or removing inconsistent landmarks in labeled data L_i_ creates a modified L_i_’ to be aligned with the unlabeled data. We score the resulting alignment by counting missing consistent adjacencies. This score is used as an objective function to drive optimization as we modify the set of inconsistent landmarks. The optimized set of landmarks ultimately yields an alignment with a maximal score, A_i_*max*. We term this process by which we optimize to pick a set of modifications, scoring them by the alignment quality co-optimization. Finally, to utilize the multiple sets of labeled data in the ensemble model, the unlabeled data is independently aligned with each labeled dataset. From the alignment with each labeled dataset: A_i_*max*, each landmark in the unlabeled data receives a predicted identity. A consensus A*max* is arrived at by a vote among the set of predicted identities for each landmark, where the most common identity is taken (Fig. 2c). The co-optimization algorithm is implemented with a correspondence-free nonlinear warping method for pre-alignment ^49^ (Fig. S3a), Linear Assignment Problem (LAP) matching ^50^, data-driven modification, and greedy optimization of labeled data (see Methods).

### Correlated Cell Identities Convey a Single-Cell View of EM Series

We apply the co-optimization algorithm to align the FM and EM series of *C. elegans* embryogenesis, using individual cells/nuclei as landmarks. We use the FM series, where cells were tracked and lineage identities assigned, to generate the ensemble model for each developmental stage in the EM series and transfer cell identities to the EM (see Methods). Due to the lack of lineaged FM data at the 2-fold stage, we only do so for the bean, comma and 1.5-fold stages in the EM series. Based on the timing of known developmental events in the FM series, the aligned EM data are timed as approximately 320, 345, and 380 min post first cell division (pfc). During this developmental window, 75 cell divisions and 76 deaths occur, while cell counts remain virtually constant (576-601) (Fig. 3a). The ensemble model for each EM dataset (Table S3) contains 39 labeled 3D FM images of embryos, each generating 5838-6170 pairwise adjacencies between landmarks, of which 439-504 (∼8%) are consistent, as well as 27 to 34 inconsistent landmarks/cells (∼5% of cells) (Fig. 3a,S4a). While the scale of the ensemble models is fairly constant over the developmental stages, the consistent adjacency graph that guides alignment is dynamic (Fig. S4a), only 146 adjacencies, about a third of consistent adjacencies, are observed at all three periods due to both inconsistent landmarks and embryo-wide cell movements during organ morphogenesis and body elongation ^31^.

**Figure 3:**
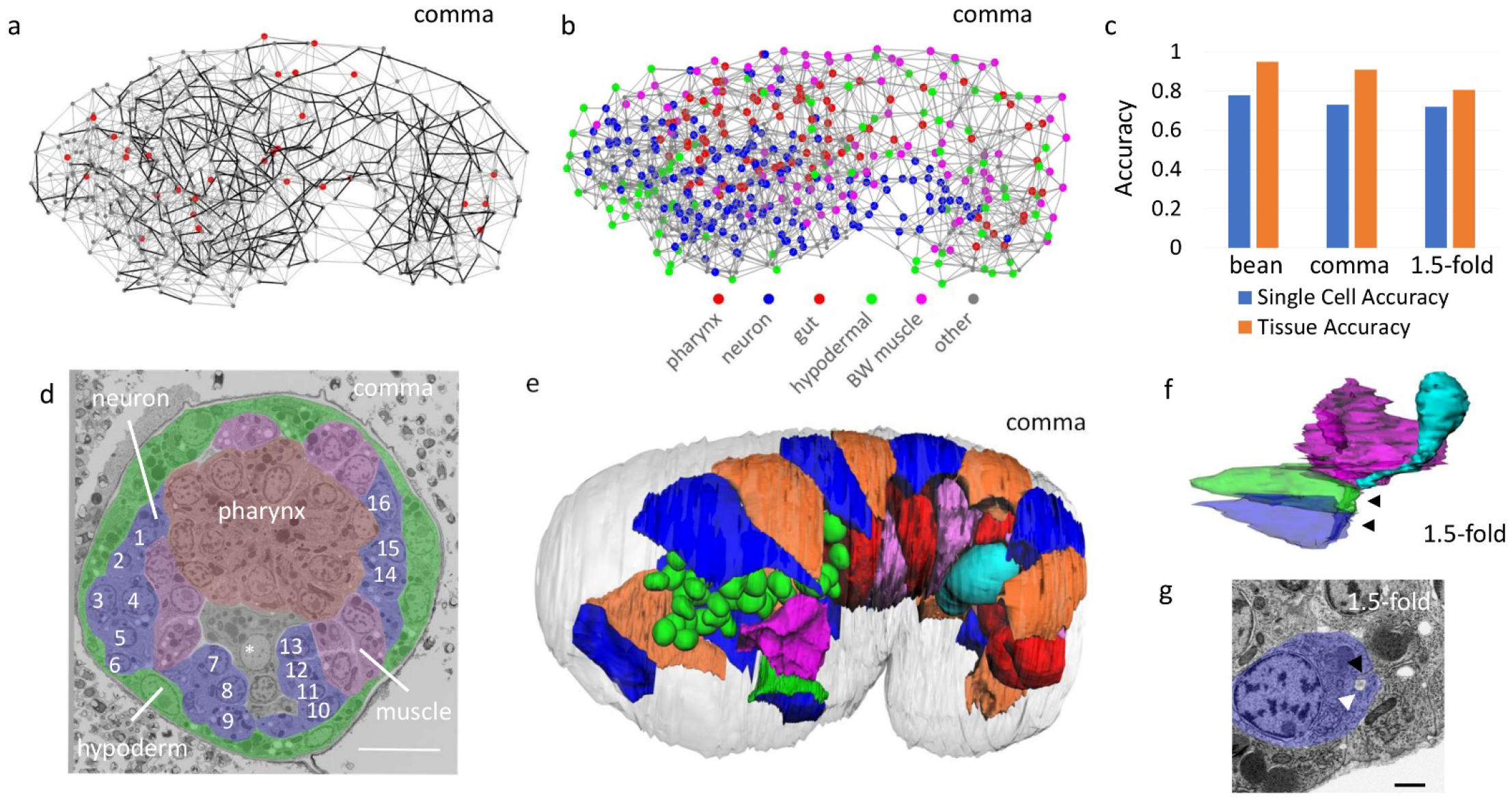
Cross-modality Alignment Conveys a Single-cell View of EM Data. **(a)** Illustration of the ensemble model for the comma stage. Landmarks (dots) and adjacencies (edges) in a single instance of labeled data. All adjacencies are shown with the consistent adjacencies highlighted in black. Inconsistent landmarks are larger and red. **(b)** A visualization of the comma stage EM data after alignment. Nucleus centroids are dots colored by predicted tissue type with color key provided. Adjacencies are shown as gray lines. **(c)** Accuracy of predicted identities for each EM data at the single-cell and tissue levels. **(d)** Illustration of EM data annotated by identities. Tissue regions are shaded following the color scheme in b. The excretory canal cell is marked with a star with the nucleus of the excretory duct and part of the pore cell body visible below it. Individual neurons are numbered. Identities as follows are from alignment (-a) or manually confirmed (-c) 1 FLPR/AIZR parent-c, 2 ASER-c, 3 AVBR-c, 4 ASHR-c, 5 AWCR-c, 6 SIBDR-c, 7 AVKR-a, 8 AIYR-a, 9 SMBDR-c, 10 SMBVL-a, 11 AIML-a, 12 AVKL-a, 13 SMDDL-c, 14 FLPL/AIZL parent-a, 15 RMGL-a, 16 ALML/BDUL parent-c. Scale bar indicates 5 μm. **(e)** 3D reconstruction in comma stage EM data showing cell contours for seam cells (blue and orange), gut cells (alternating shades of red), germ line (cyan), and the excretory canal, duct and pore cells (red, green, and blue respectively) as well as nuclear contours for pharynx (green). **(f)** 3D reconstruction of the excretory system in the 1.5-fold stage EM data. Red, green, blue and cyan are the excretory canal, duct, pore and gland cell, respectively, as in e. Black arrows point to auto fusion in the duct and pore cell. **(g)** EM view of lumen (white arrow) and site of auto fusion (black arrow) in the excretory pore cell at 1.5-fold stage. Scale bar indicates 1μm.

The result is a single-cell level alignment between the FM and EM series, with a predicted identity for each cell in the EM series (Fig. 3b, S4b), which we refer to as the correlated EM series. We assess the accuracy of the correlated cell identities based on a set of manually curated cell identities across organs and tissues (Fig. S4b, 148 to 242 per embryo). We consider accuracy at two levels: individual cells and tissues (Fig. 3c). Single cell accuracy, which is the agreement between manual and predicted identity, varies between 71-78%, while accuracy of tissue identities is 79-95%, both with a trend for decreasing accuracy for the later developmental stages. Among head neurons, the single cell accuracy is 69-78%. There is no comparable embryonic work, but this is higher than previous reported efforts that attempted to align and name all head neurons in young adults based purely on cell positions (50-68%) ^17^.

These results are deposited on webKnossos ^51^ as a community resource containing the EM images, annotated nuclear positions and cell identities. Image data and identities are navigable within a convenient interface which allows users to fork and revise annotation. Notably, 56-70% of the cell identities are manually curated (Fig. S4b), including 25-40% as cell identities and 16-34% as tissue/organ identities, greatly reducing potential errors in identities. Annotation will be continually updated based on community feedback. The correlated cell identities convey a single-cell view of the EM data for developmental dynamics at different scales (Fig. 3d,e), such as tube formation in the excretory system through concerted auto fusion of cells to connect the canal to the outside environment (Fig. 3f,g) ^52^.

### Spatial and Temporal Organization of Neuropil Formation

Leveraging the co-optimization algorithm for alignment, we conduct cross-modality analysis of early NR development to elucidate the spatial and temporal organization underlying neuropil formation. The nerve ring (NR) is the primary neuropil in *C. elegans*. The adult NR consists of 181 neurites, where neurite topography and contacts constrain synapse formation and circuit function ^27-29,53^. The NR has long been a prime model to study the general principles of brain structure and function. The NR emerges in the later half of embryogenesis ^54^, but our understanding of the spatial and temporal organization of neurite outgrowths in this process is still poor. Efforts with FM revealed that pioneer neurites appear around the bean stage and a ring around the pharynx becomes visible by the comma stage (Fig. 4a)^29^. FM with cell-specific markers and sparse labeling has led to critical insights: identification of pioneers ^29,34,36^, the inside-out pattern of organization of outgrowth ^29^, and even neurite topography through combinations of two-color imaging (Fig. 4b). However, lack of markers to systematically label the entire nervous system and lack of spatial resolution hinder further insights. EM on the other hand provides a global view of not only every neurite but also the surrounding tissues with high spatial resolution (Fig. 4b). We scan through EM data to identify neurites, and back trace them to soma in order to obtain identities. We also correlate EM with a set of FM series that encompasses 12 strains with cell specific markers that label neurons individually or in small, isolated groups (Fig. S1, Table S4) ^29,46^.

**Figure 4:**
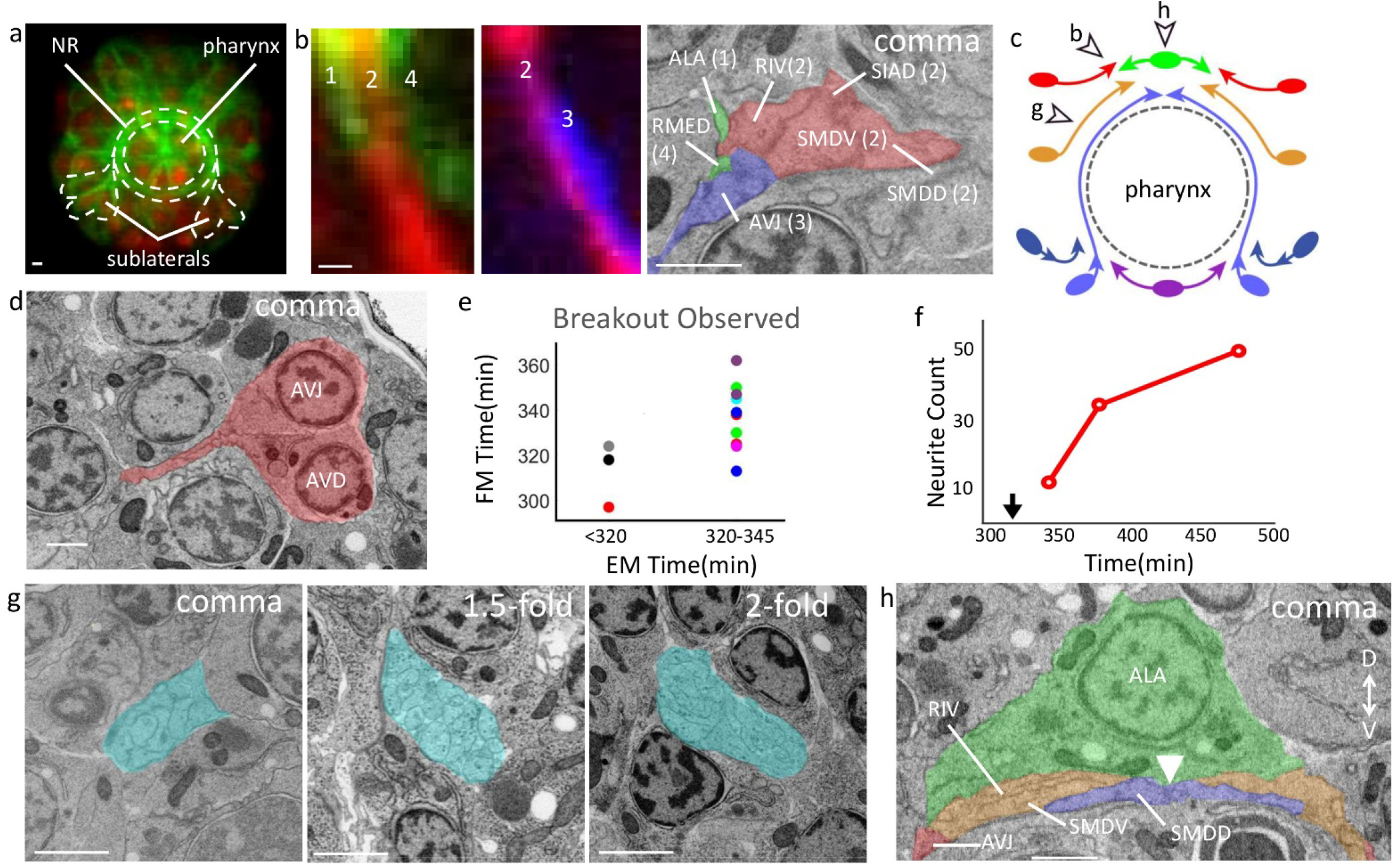
Spatial and Temporal Organization of Neuropil Formation. All scale bars 1 μm. **(a)** FM cross section of a comma stage embryo with a ubiquitous histone marker (red) and a broadly expressed cell membrane label (green) ^31^. Anterior view. Dotted line encircles the NR and routing of sublateral neurons into the ring. **(b)** Comparison of resolution in resolving NR in FM and EM. Two images of two-color FM image of developing neurites (red, green and blue are labeled by the promoters of *cnd*-1, *ceh*-10 and *zag*-1, respectively). Neurite bundles are numbered anterior to posterior. EM showing the corresponding neurites and their placement in the ring at the comma stage. Neurites are labeled with names and colors of the corresponding bundle in FM. **(c)** Schematic of the NR (anterior view) showing pioneers, entry points, and approximate extent at bean and comma stages. Ovals represent neuronal soma and arrowheads the leading edge. Two arrowheads are shown for the amphid (dark blue) and sublateral (light blue) outgrowths to demarcate progress at bean and comma stages. Moving dorsal to ventral, dorsal outgrowth is green, supralateral red, lateral orange, and ventral purple. See Fig. S3, S4 for corresponding EM. Open arrows and associated letters indicate the figure panels that show the location pointed to. **(d)** EM at comma stage showing the pioneers of the supralateral entry point with cells shaded red and named. **(e)** Comparison of breakout timing derived from FM and EM. Each dot represents a single neuron or small cluster of neurons. Dot color indicates source marker in FM (*lim*-4:red, *unc*-86:green, *zag-1*:*blue, ttx-3:magenta, egl-13:cyan, cnd-1:black, ceh-37:gray, ceh-10*:purple). **(f)** Number of neurites visible in cross sections in g across the EM series. Black arrow indicates bean stage. **(g)** Axial NR cross sections (shaded cyan) near the lateral midline entry point (arrow g in panel c). **(h)** Precision in neurite outgrowth dynamics. A resliced view of the dorsal midline at comma stage shows the point left and right sublateral pioneers meet (white arrow). Colors match schematic in c.

Our analysis shows that the establishment of the NR is organized by eight entry points which include the dorsal midline, the ventral midline, and three left/right symmetric pairs in between, namely supralateral, lateral and sublateral (Fig. 4c). The early time points in the correlated EM series reveal the pioneers that establish each entry point (Fig. 4c, 4d, S5 and S6). Sublateral neurons, namely SIAD and SMDD, are the first neurons to initiate outgrowth, with the initial breakout captured at the bean stage (Fig. S5b). This observation is consistent with these neurons serving as early pioneers for the NR ^29,34,36^. In less than a half hour after this breakout (comma stage), all entry points have been populated with the corresponding pioneer neurites. Moving from the dorsal midline to the ventral midline: at the dorsal midline, RMED and ALA have grown bilaterally (Fig. S6b); supralateral neurons, namely AVD, AVH, and AVJ have entered and grown dorsally (Fig. 4d); lateral neurons, namely SMDV and RIV have also entered the ring almost reaching the dorsal midline (Fig. S6d); and at the ventral midline, RIR, RMEV and RIH have bilaterally advanced dorsally (Fig. S6g). While the EM series provide a comprehensive and definitive set of entry points and pioneers due to its label-free nature, the FM series provide much finer temporal resolution of outgrowth dynamics. In terms of the timing of neurite breakout (Fig. 4e, Table S4), breakouts observed as having occurred in EM at the bean stage span almost 30 minutes in the FM while breakouts observed in EM at the comma stage span an hour. Most of the FM times are congruent with the upper bound of timing provided by EM. Further data would clarify if the occasional contradiction in timing is due to biological variability or technical issues in temporal alignment.

In addition to defining the pioneers and the initial breakout times, the cross-modality analysis also reveals dynamics of subsequent neurite growth in the NR (Fig. 4f). Assayed near the lateral entry point (Fig. 4g, arrow g in Fig. 4c), the number of processes visible in the NR increase rapidly from 12 at the comma stage to 51 by the 2-fold stage. Outgrowth is not reducible to the timing of breakout, subsequent extension also has striking dynamics. For example, pioneers of the amphid commissure, namely ASH, RIB, and AWC, show initial outgrowth at the bean stage, which make them one of the first to breakout in the entire nervous system along with the sublateral neurons ^54^ (Fig. S5c). However, these neurites, and additional followers from the amphid neurons pause soon after their breakout ^29,46^ near the sublateral entry point (Fig. 4c, S6e) before they enter the NR (by the 1.5-fold stage).

Finally, the spatial resolution of the EM series highlights the precision of control in neurite outgrowth kinetics. For example, comparing the left- and right-side behavior of the first cells to grow into the NR, namely SIADL/R and SMDDL/R, suggests strong symmetry in behavior. Both sides have broken out at the bean stage, with similar sized outgrowths. The contralateral pairs then reach the midline simultaneously at the comma stage: their tips meet precisely beneath the midpoint of the ALA nucleus, which marks the dorsal midline (Fig. 4h). This level of precision is not limited to the sublateral pioneers. Left and right counterparts among the lateral and supralateral pioneers have also reached nearly identical positions relative to the dorsal midline at the comma stage (Fig. 4h). It is not clear how such level of precision is achieved, or if there is signaling to coordinate between the two sides.

### Cell-Cell Interactions and Ultrastructure During Amphid Organogenesis

The amphid is a major sensory organ of *C. elegans* consisting of 12 neurons and two glial cells (the socket and sheath cells). The adult sensory end is a bundle of ciliated dendrite endings with drastically different morphologies surrounded by a tube-shaped glial channel. The glial cell extensions are in turn embedded between skin (hypoderm) cells to create a sensory structure that is open to the environment ^54,55^. The dendrites grow via a distinctive mechanism, a collective retrograde extension from an initial rosette structure towed by the migrating hypoderm ^32,33,56^. Formation of the embedded opening and elaboration of dendrite morphology occur in embryogenesis but are less understood ^55^.

The correlated EM series confirms the dynamic cell behaviors revealed by FM during dendrite extension (Fig. 5a) ^32^ and adds structural details. For example, 3D reconstruction at the bean stage shows small protrusions from amphid neurons clustered as in the initial rosette and lodged in a dent at the anterior/leading edge of the hypoderm (Fig. 5b,c, S7a,b,c). Notably, the socket cell, though engaged in the rosette earlier does not appear engaged in this focal location at the bean stage and only touches a handful of amphid neurons (Fig. 5b, S7a,b).

**Figure 5:**
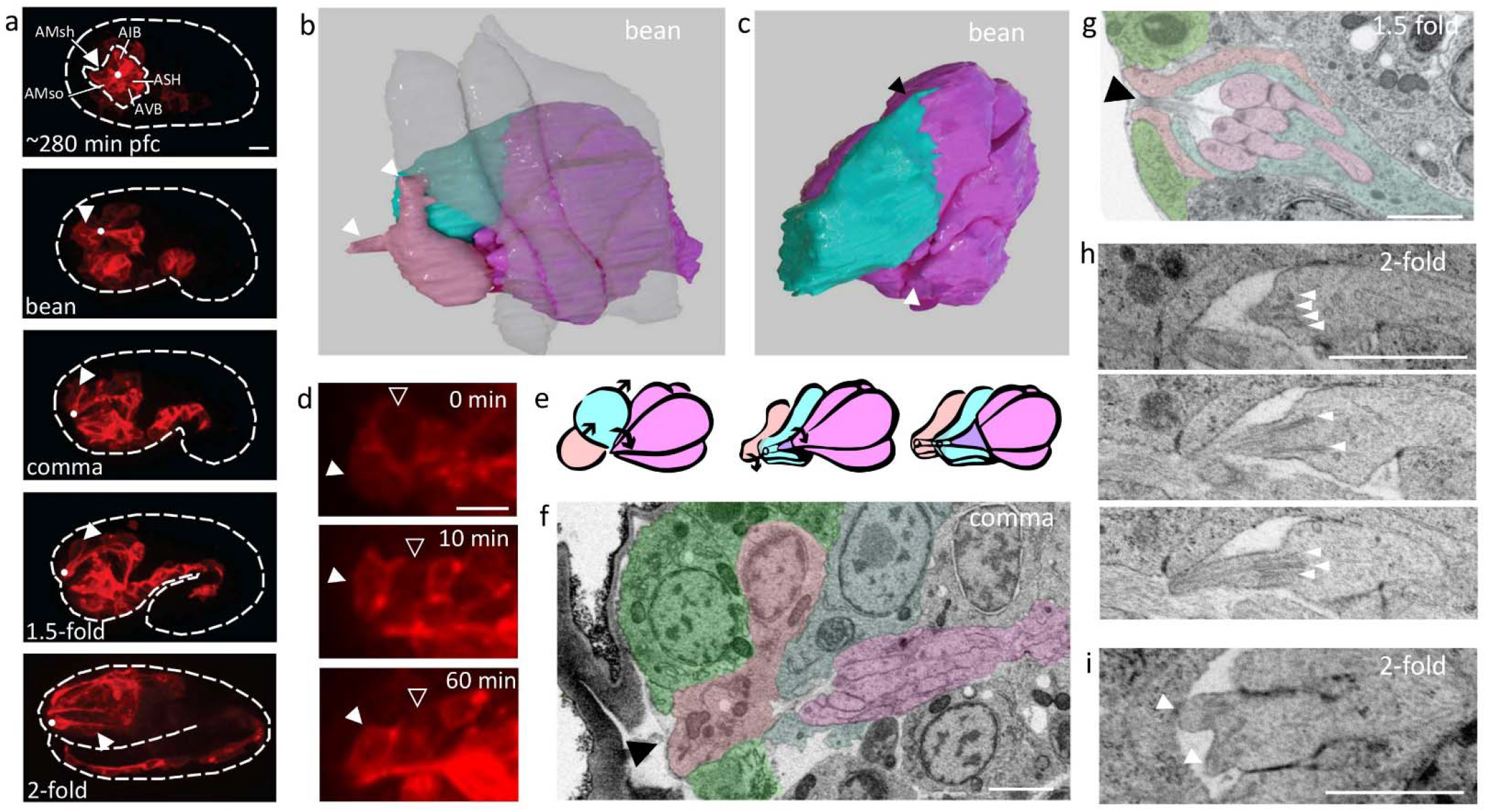
Cell-Cell Interactions and Ultrastructure During Amphid Organogenesis. All scale bars 1 μm. **(a)** FM view of amphid dendrite development with a *cnd-1* promoter driven membrane label. Dashed line shows embryo contour. Dot indicates the location of dendrite tips, and arrow the socket and sheath cells. Dashed circle inside the embryo outlines the initial amphid rosette at ∼280 min pfc. **(b)** 3D reconstruction of cells participating in the collective dendrite extension from bean stage EM. Hypoderm: semi-transparent; amphid neurons: magenta; sheath cell: cyan; socket cell: pink. Arrows indicate protrusions on the socket cell. **(c)** The same reconstruction as in b with socket and hypodermal cells removed. Black arrow highlights protrusion on the sheath cell. White arrow indicates focal point of amphid neuron protrusions. See also Fig. S6. **(d)** Partial max projection of FM image with a *cnd-1* promoter driven membrane label showing socket cell (filled arrow) and sheath cell (open arrow) posterior migration. **(e)** Schematic of the time course of sensory opening formation. Arrows indicate motion of cells. Colors match b. **(f)** Resliced view of EM at comma stage highlighting socket cell (tinted rose) shape and relative position to hypoderm cells (green), sheath cell (cyan) and dendrites (magenta). Black arrow indicates contact between the socket cell and the exterior of the embryo. **(g)** EM at the 1.5-fold stage showing the sensory opening. Colors follow f. Arrow indicates the opening to the exterior of the embryo. **(h)** EM at 2-fold stage showing three sections of a dendrite tip. White arrows mark nine individual microtubule structures. **(i)** EM at 2-fold stage showing a bifurcated dendrite tip, each tip marked with an arrow.

More importantly, our cross-modality analysis reveals critical details of cell behaviors and interactions in creating the multi-layered structure of the open sensory end. The sheath cell starts on the anterior side of the amphid rosette (Fig. 5a). Subsequently, the sheath cell body moves dorsal-posteriorly (Fig. 5d). Lamellipodia-like protrusions from the sheath cell during this process are visible at the bean stage (Fig. 5c). Meanwhile, the anterior portion of the sheath cell wraps itself around the collected mass of the dendrite tips (Fig. 5e), which is evident by the comma stage (Fig. 5f). The socket cell exhibits more complex behaviors. One set of behaviors follows a similar course as the sheath cell. The cell body migrates posteriorly (Fig. 5d), creating transient posterior protrusions (Fig. S7d), while the anterior end remains close to the dendrite tips. Subsequently, after the sheath cell has wrapped the dendrites, the socket cell starts to wrap over the sheath cell (Fig. 5e,f). The wrapping is complete by the 1.5-fold stage (Fig. 5g) ^33^. Distinctively, the socket cell must also interface with the hypoderm to create the open sensory end (Fig. 5g). Noteworthy details hint at the dynamic role of the socket cell in the process. During dendrite extension, the socket cell makes elongated protrusions, one of which appears to reach out and contact the advancing hypoderm cell (Fig. 5b). Later, the anterior end of the socket cell is juxtaposed between hypoderm cells and exposed to the outside of the embryo like a plug (Fig. 5f), which may play a role in creating the opening to the environment in the sensory end. The socket cell is fully surrounded by the hypoderm by the 1.5-fold stage (Fig. 5g, S7e). Overall, the dynamic details from the cross-modality analysis suggest that the strategy of amphid sensory end assembly is successive wrapping of interior structures with additional layers.

Furthermore, the series illustrates the elaboration of distinctive ultrastructure. Centrioles move to the dendrite tips between the bean and comma stages (Fig. S7f) ^57^. By the 2-fold stage, centrioles have disappeared, presumably degraded ^57^, and microtubule bundles (9 bundles per dendrite) are visible growing into the short proto-dendrite tips (Fig. 5h). Their appearance is essentially the same, aside from length, as adult structures ^55^. Meanwhile, the dendrite tips begin to elaborate their complex morphology. The shape of the dendrite tips remains simple up to the 1.5-fold stage. By the 2-fold stage, some diversity in length and shape is visible. Notably, two dendrites, namely ADF and ADL, show short bifurcated tips, each with its own set of microtubules branching from a central location ^55^ (Fig. 5i). Dendrites tips are not yet individually embedded within the sheath cell by the 2-fold stage.

## Discussion

Our study demonstrates a powerful co-optimization algorithm to align and assign identity to landmarks. Because each cell in *C. elegans* is unique, it is often desirable to identify individual cells. For postembryonic images, people combine cell position with the characteristic cell/neurite morphology to make a positive identification. In the embryo, before such morphology becomes prominent, cell identification has been challenging. In his seminal work, Sulston suggested that cells in the late embryo can be identified by careful examination of relative positions within small neighborhoods ^44^, but he remained the only person who could do so systematically. Instead, researchers use live imaging-based lineage tracing for reliable cell identification ^42,58^. By paying attention to relative position through the adjacency graph, our algorithm essentially follows Sulston’s suggestion, and achieves unprecedented accuracy in cell identification. Additional information can be readily added to the adjacency graph to better score matched landmarks and improve the alignment, such as size, appearance and other features of the landmarks when available. Furthermore, while our study is focused on the wild type, one could use live FM to build the appropriate ensemble models to analyze mutants. Thus, by eliminating the bottleneck in data acquisition as well as data interpretation, we envision a broad use of EM and cross-modality analysis of *C. elegans* for both the wild type and mutants. As a proof of concept, we present the image data and identity annotation of our correlated EM series as an accessible public resource. Existing annotation provides a navigational aid and the seed of a community effort that we hope will fully validate annotation of the series.

More broadly, the co-optimization algorithm provides a general solution for cross-modality and cross-scale alignment. Representation of landmarks as a point cloud is modality- and scale-free. In our study, we use individual cells/nuclei as landmarks due to the desired single-cell resolution for *C. elegans* biology, but the point cloud can easily represent a mixture of landmarks at different scales, enabling ambitious future work. With advances in deep learning and image analysis, landmarks can be extracted from complex organs such as the mammalian brain by learned region detectors ^12^ with increasing reliability. While the mammalian brain is orders of magnitude larger in terms of cells, it is worth noting that in state of the art atlases it is currently annotated with hundreds of regions ^14^, which makes alignment at this level of detail a similar scale problem to ours. We postulate that at an optimal level of landmark density consistent adjacencies provide an effective implicit model of anatomy that can maximize available prior information about structure and constrain accurate automated alignment. The data-driven modification in the co-optimization algorithm further facilitates such a vision for automated alignment and annotation of complex brain images. It readily handles not only biological variability in landmarks (developmental or between individuals) but also technical variability such as errors in landmark extraction or systematic differences between modalities. This model only becomes more powerful as data accumulates. Ultimately, we imagine a multi-modal fusion process built on our approach will be capable of discovering consistent landmarks bottom up and systematically collating information from multiple modalities. This would automate the decision making involved in defining the regions of a digital atlas ^14^ making a full spatio-temporal atlas of developing anatomy possible.

Last but not least, our post-hoc correlation of EM and FM data provides a useful alternative to true correlative EM. Correlative EM is a powerful, but complex to implement, approach for targeted EM based on the expression pattern of fluorescence markers ^59^. We note that when used with Array Tomography, the identity of landmarks obtained from our post-hoc correlation can be used to select targets for re-imaging with EM at higher resolution.

## Methods

### 1. EM Imaging

#### 1.1 High-pressure freezing and freeze-substitution

A generous amount of *E. coli* was placed into 0.5 μm carriers filled with 20% BSA solution. Embryos from N2 worms were collected using a platinum pick from the seeded plates and settled in the carrier. The carriers were covered with a lid and frozen in a Wohlwend HPF Compact 02 (Wohlwend GmbH, Sennwald, Switzerland).

The freeze-substitution step was done in the freeze-substitution unit (AFS2; Leica Mikrosysteme GmbH, Vienna, Austria) following the previously described procedure ^60-62^. Briefly, frozen samples were transferred under liquid nitrogen to the freeze-substitution mix of 2% (w/v) osmium tetroxide, 0.1% (w/v), uranyl acetate 2% (v/v) H2O in acetone in acetone pre-cooled to -120°C. Samples were heated to -90°C over the period of 2h and left at this temperature for 2h. Over 24h, the temperature was raised to -30°C. Samples were left in this temperature for 2h. After they were washed once with acetone and three times with ethanol, the temperature was raised to +20°C. The solvent was replaced with the mix of ethanol and Epon resin mix (Embed 812 resin (EMS, 1420), infiltrating the samples with increasing concentrations 30%, 50%, 75% for three hours each exchange at ambient temperature. Three incubations and exchanges of 100% EPON were done over 12h. Samples were flat embedded using the two-step procedure ^62^ and polymerized for 48h at 60°C.

#### 1.2 Block trimming

For accurate sectioning it is important to orient samples correctly. Flat blocks facilitate recognition of a ROI and help in trimming. Delimited areas surrounding the sample were trimmed using a 90 diamond trim tool (Diatome, Biel, Switzerland). Care was taken to approach the sample as closely as possible and keep the top and bottom sides parallel to one another and the knife’s surface ^63^. A buffer area of empty resin at the leasing edge of the trimmed sample was left to allow space for alignment maneuvers during the subsequent steps.

#### 1.3 Focused ion beam/scanning electron microscopy (FIB-SEM)

For FIB experiments, samples were oriented transversely or longitudinally already at the stage of trimming. Sample blocks with targeted trimmed rectangles ∼100×200 μm were glued on aluminum SEM specimen stubs and coated with a 30 nm layer of platinum. Stubs were introduced into FIB-SEM (Helios Nanolab 650, Thermo Fischer Scientific) and further calculations were done inside the microscope chamber to target the ROI precisely to the embryo of interest. FIB/SEM acquisitions were performed using FEI Slice and View software™. The FIB milling was done at 30 kV acceleration voltage, 790 pA current. Images were acquired with the in column detector (ICD, comma stage embryo) and mirror detector (MD, bean stage embryo) in immersion mode using an electron beam of 2 kV, 800 pA and 10 μs of dwell time.

#### 1.4 Array Tomography (AT)

For AT experiments, the “blind” sectioning strategy does not require precise sample orientation. Several samples can be sectioned at once, and assessment of age is done after sectioning.

Planar arrays of the section were generated using the ATS knife (Diatome, Biel, Switzerland) mounted on a Leica UC7 ultramicrotome (Leica Microsystems; Vienna, Austria). Rectangles of silicon wafer approximately 2×4 cm (Ted Pella, No 16015) were cut from the round plate glow discharged, cleaned, and placed into the knife boat. “Ribbons” of ∼300 consecutive sections, 70-100 nm thick, were subdivided into shorter segments and then aligned side by side on support ^63^. After the sections transfer and water drains, the wafer was let dry slowly at the ambient temperature and subsequently incubated for 30 min at 60C.

The wafers were transferred to the Helios 650 FIB-SEM (Thermo Fischer Scientific). The strategy of positional correlation to select the embryos of the desired stage was used. To find the anchor sections that are worthy of being analyzed in sequence, we chose the strategy of “leaping” between centrally located sections across the short horizontally aligned “ribbons” to screen for the ROI. Acquisition parameters were similar to those used in FIB protocols. Images were acquired with mirror detector (MD) in immersion mode using an electron beam of 2 kV, 800 pA and dwell time of 5-10 μs.

#### 1.5 Image analysis and rendering

For alignment of slices within an EM data set, we used IMOD or FIJI. Final stacks were contrast inversed to achieve a TEM-like appearance. 3D tracing of sections was done with the 3dmod module of IMOD^64^ or TRACKEM^65^ with 3D rendering done in Daz3D ^66^.

### 2. 3D, Time-Lapse FM Datasets

As previously described ^29^ time lapse images of fluorescently labeled *C. elegans* embryos were collected with spinning disc or diSPIM microscopy ^67^, and lineaged with StarryNite ^42,47,68^ to assign cell identities. The annotated nuclear centers serve as landmarks for ensemble model creation and series correlation.

### 3. Cross-Modality Alignment

#### 3.1 Pre-alignment

Pre-alignment between labeled and unlabeled data allows the use of absolute spatial position and is performed in two steps.

First, an approximate affine least squares or Thin Plate Spline (TPS) ^69^ alignment is performed based on a small number of manually identified landmarks. The landmarks are identified in each EM dataset based on their characteristic positions and cell shapes. Cells from different regions of the embryo are chosen to highlight the body axes (Table S5). The landmark set was evaluated visually per EM data set to ensure the point clouds of labeled and unlabeled data fully overlapped when aligned.

Second, the approximate alignment is refined using a coherent non-linear method GMM-CPD ^49^. This method creates a regularized local warping function to align two point clouds by iteratively warping and matching points under an assumption that nearby points move similarly, which provides a more granular adjustment for non-linear shifts in body proportion and relative positioning of organ structures (Fig. S2a).

#### 3.2 Temporal Staging

A coarse temporal staging is used to select the appropriate time window in the FM data to create the ensemble model. Stage is visually estimated based on matching body axis elongation in FM and EM datasets, followed by the pre-alignment as described above. A visualization of manual correspondences under pre-alignment is then used to adjust the temporal staging if alignment results appear to contain a systematic AP shift related to the elongating body axis.

We note that quantitative scores from the alignment method described below can be used for precise temporal alignment (Fig. S2b) which would be useful in implementing fully automated methods of spatio-temporal alignment. A wide and smooth minima around the manual temporal alignment demonstrates a minimization of constraint violations over different possible temporal alignments would provide a similar temporal alignment to our manual assessment. The absolute minima being offset by a few minutes from our manually assessed staging suggests the approach might even improve on manual time estimate. However, such a precise temporal alignment is not needed for the purpose of choosing a relatively wide time window to create the ensemble model.

#### 3.3 Ensemble Model Creation

For each EM stage an ensemble model encompassing adjacency, position and inconsistent landmark information is created to be used in alignment. Adjacency and positional information is collected within a local (13 min) window of time in each of the three FM embryos that contribute to the labeled data ensemble. Temporal window size was optimized to maximize identity prediction performance using one of the three labeled data embryos as a test case and the other two as labeled data. Adjacencies are predicted via a Gabriel graph^70^ for all landmarks in a labeled dataset. Consistent adjacencies are those present in all datasets within the ensemble. A list of inconsistent landmarks is compiled from all cell deaths, and cell divisions that occur within the local temporal window.

#### 3.4 Modification and Co-optimization

We use our ensemble model to guide modifications of inconsistent landmarks in labeled data optimizing the exact set of labeled data landmarks along with alignment. This optimization uses missing consistent adjacencies in the labeling of the unlabeled data which is created by the alignment as an objective function in a process we term co-optimization.

Each possible modification to the labeled data represented by the list of inconsistent landmarks is tested. The hypothesis that a programmed cell death has already disappeared in the unlabeled sample is considered by removing that cell from the labeled data. For divisions the hypothesis that a recently divided cell in the labeled data has not divided in the unlabeled sample is considered by merging the two sibling cells into one parent cell at their midpoint. The hypothesis that a cell in the labeled data that will soon divide has already done so in the unlabeled dataset is entertained by splitting the parent cell into two cells sharing the same location. Note that while it would be possible to use prior information about the expected division axis to create more accurate daughter positions, this seems over complicated as the main goal is to prevent cascading errors due to qualitative discrepancies in the set of landmarks present.

A simple, greedy, gradient descent process optimizes the labeled data. The optimization first considers the (limited in *C. elegans*) possible divisions and then deaths and accumulates changes that improve total adjacency constraint violations. For each of these hypotheses a match of the unlabeled dataset against the modified labeled data (L_i_’) is made and adjacency constraint violations are counted under the corresponding alignment (A_i_). A proposed change is kept if it results in a significant (in our case>=3) reduction in adjacency violations.

#### 3.5 Linear Assignment and Scoring via Consistent Adjacencies

For a given pre-aligned unlabeled dataset and modified labeled dataset, matching is framed as a linear assignment problem (LAP) minimizing the total distance between pairs of points. Missing consistent adjacencies are then used to score these matching results to evaluate modifications and to identify and prohibit incorrect matches.

Given an alignment between a modified version of the labeled data and unlabeled data that alignment can be evaluated by computing the Gabriel graph for the unlabeled data and, using the predicted identities under the alignment, counting missing consistent adjacencies. The total number of missing consistent adjacencies for the alignment is used to score the modification process described above.

In addition, a high number of missing consistent adjacencies for an individual landmark’s predicted identity (summed over all expected adjacencies of that landmark only) indicates it is likely wrong (note this can be assessed only given the global alignment, this cannot be assessed independently for a single landmark match). These individual match scores are used to identify and attempt to correct incorrect matches in an iterative process. The individual identity assignment with the worst score, (i.e. the most missing consistent adjacencies) if any exists, is made impossible (set to infinite cost) and LAP matching is repeated. This process is repeated for N iterations, and at the end of those iterations the global alignment with the lowest total violations is selected. Iterative error correction (200 steps) is performed after each modification, before judging its quality. Once a labeled dataset is fully co-optimized an additional 1400 iterations are performed and the best alignment and matching for this is A_i_*max*.

#### 3.6 Voting and Reiteration

Voting picks an identity for each landmark from the set of predicted identities corresponding to an alignment of the unlabeled data against each labeled data set. The most common name out of these independent answers for each landmark in the unlabeled data is taken as its identity. This procedure can result in multiple cells in the unlabeled data being assigned the same identity. Though it would be possible to force a unique set of identities by using an additional linear assignment between landmarks and available identities scored by total votes for each identity per landmark, we do not perform this step as we found this marginally reduces accuracy (data not shown).

Highly reliable landmark names can be used to reinitialize the alignment more accurately improving matching results. Landmarks in the sample where all or nearly all assignments agree are most reliable, (see Fig. S2c). Taking landmarks where the winning identity has a 75% supermajority in place of the TGMM automated nonlinear alignment we compute a new landmark based TPS alignment based only on these ‘known good’ matches. These reliable automatic matches are greater in number than initial landmarks and can potentially provide a better nonlinear pre-alignment than TGMM. We then repeat the exact same co-optimization based matching for only the subset of uncertain cases, (the certain cases are reinserted with their fixed identities when evaluating adjacency constraints). All reported results include one round of iterative re-alignment.

#### 3.7 Algorithmic Ablation Results

To evaluate the contributions of distinct components of our method to performance we compare performance after ablating parts of the algorithm to our full algorithms’ performance (71-78%). Ablating all of the unique components of our approach leaves the prior method of a LAP matching based on distance (after pre-alignment) to each labeled example followed by voting, this has accuracy of 43-66%. The most novel component of our system the co-optimization of unstable landmarks makes a significant contribution to accuracy; if we skip this step single cell accuracy ranges from 67-75%.

#### 3.8 Computational Considerations

Our implementation has a significant runtime but could be significantly optimized if desired. Our co-optimization algorithm was implemented in Matlab. Alignment was performed on a Xeon Gold 6128 3.4GHz CPU with 500mb of memory (alignment is however not particularly memory intensive). Runtime was variable depending on the temporal stage and number of modifications to be considered but took about a day. The vast majority of run time is taken by the tens of thousands of iterations of linear assignment implicit in the modification and constraint optimization process. Results are fairly insensitive to the number of constraint iterations used. Reducing iterations and total computation time by 3/4 results in only a small decrease in accuracy. Given the small number of datasets computation is not a major issue and reported timing and accuracy results reflect a large number of iterations run after initial landmarks and parameters were optimized using runs with a low number of iterations. A more efficient approximate LAP implementation would significantly decrease computational costs as over 95% of computational time is spent in this function. Runtime would also be significantly reduced by an explicitly parallelized implementation as alignment of individual labeled datasets, currently sequential, is completely independent; this was not attempted.

#### 3.9 Manual Curation of Cell Identities

A significant number of cells in each EM dataset were manually curated to establish their identities at the tissue or individual cell level. Tissue type could be easily assessed for pharynx, hypoderm, gut, and body wall muscle from morphology for most cells at all stages. To identify specific individual cells we started at cells with distinctive morphologies and then distinctive positions on the exterior of the embryo or adjacent to the pharynx. A subset of these are landmarks used in initial pre-alignment. Moving out from these highly reliable landmarks we examine labeled and unlabeled data sets as 3D point clouds and identify cells with consistent position relative to a landmark in labeled data sets and the corresponding cell in EM. These identifications are further checked against EM morphology, and known neuronal appearance from FM. Identified patches are expanded checking for inconsistencies until all cells in a small local patch are identified. Total time spent on annotation varied between a few days for the least annotated 1.5-fold data set and a bit more than a week for the best annotated comma data set.

## Supporting information

EM cell annotation bean

EM cell annotation comma

EM cell annotation 1.5-fold

## Acknowledgements

Thanks to Bruno Humbel, Jean Daraspe, Helmut Gnaegi, Tilman Franke for advice and assistance with EM, Chris Brittin, Li Fan, Kris Barnes and all the members of the WormGUIDES consortium for advice regarding *C. elegans* neurodevelopment. Particular thanks to Chris Brittin, Daniel Colón-Ramos, Hari Shroff, Mei Zhen, and Ryan Christensen for advice and comments on the manuscript. This study was partly supported by NIH grants to Z. B. (R01GM097576, R24OD016474). Research in Z.B. lab is also supported by an NIH center grant to MSKCC (P30CA008748). A.S. was supported by grant 2019-198110 (5022) from the Chan Zuckerberg Initiative and the Silicon Valley Community Foundation.

## Author contributions

Conceived algorithms: AS. Designed and conducted EM imaging: IK, CK. Drafted manuscript: AS, ZB. All authors provided critical revision of manuscript. Supervised research: IK, ZB.

## Data and materials availability

EM data and annotations available on webKnossos from project site: https://sites.google.com/view/developmentalem/home

Alignment code is available on Github: https://github.com/zhirongbaolab/CellIDAlignment

Additional, correspondence and requests for materials should be addressed to I.K. and Z.B.

