## Supplemental Figures and Tables for "Cross-modality Synthesis of EM Time Series and Live Fluorescence Imaging"

**Supplemental Materials**

**Supplemental Figure Captions:**

**Figure S1:** Overview of FM datasets. Max projection of representative time point for the FM data used. Each strain carries a ubiquitous histone marker (red) and a cell membrane marker (green) driven by a specific promoter (shown in each picture). See also Table S2.

**Figure S2:** Pseudo-Code of Algorithm. Outline of steps of algorithm giving an explicit sense of how various model components and sub algorithms fit together in our approach.

**Figure S3:** Method Details. **(a)** Pre-alignment Example. Effect of nonlinear pre-alignment. Equivalent nucleus locations in two temporally aligned FM data sets are connected with a tapered cone to show displacement. The top image is these displacements after an optimal affine alignment calculated from all cells. Significant displacement between individuals cannot be accounted for by a global affine alignment. The second image shows the same displacements after applying a regularized nonlinear alignment method that does not assume known correspondences GMM-CPD on top of this initial affine alignment. **(b)** Plot of missing consistent adjacencies for alignments from our approach between the bean EM data and one FM embryo. The labeled data set and expectation model used is set to various temporal offsets from the manually derived optimal temporal alignment. The minimum is centered on the manual alignment, indicating an automated search for an optimally scoring temporal alignment would have produced a similar result. The precise minima is not identical suggesting automation could be used to refine alignment. **(c)** Consistency of identity votes as a quality metric. ROC chart for the fraction of total correct and incorrect identities returned as we accept identities from our alignment of the bean and comma state with fewer votes for the winning identity. This is compared to a ubiquitous alternative quality metric on the same results: distance between matches (computed using one arbitrarily selected prealigned FM data set). For much of its length the votes curve is to the upper left of the distance curve showing it returns more correct answers for a given fraction of incorrect answers.

**Figure S4:** Cross-modality Alignments. **(a)** Illustration of the ensemble model for each EM dataset aligned. See Fig. 3a for a detailed explanation. **(b)** A visualization of all named EM data. Nucleus centroids are dots colored by predicted tissue type, matching the color key in panel b. Computed adjacencies are shown as gray edges. **(c)** Summary of annotations of each EM dataset. A histogram stratified by tissue type for the type of identity available for each cell. Manually verified single cell or tissue type IDs are in blue and red respectively. The identities predicted by our alignment method are gray.

**Figure S5:** Earliest Neurite Outgrowths. **(a)** Illustration of cell groups at bean stage and approximate extent as in Fig. 4c. **(b-c)** EM image of each breakout location with cell areas shaded to match cartoon. End of neurite or end of cell body from which the neurite is growing is indicated with an arrow.

**Figure S6:** Pioneer and Other Early Neurites at the comma stage. **(a)** Illustration of cell groups at comma stage and approximate extent as in Fig. 4c. **(b-g)** EM image of each location with cell areas shaded to match cartoon. End of neurite or end of cell body from which the neurite is growing is indicated with an arrow. In e this also indicates the point where amphid cells pause in their outgrowth.

**Figure S7:** Amphid Organogenesis. **(a)** Cross section of dendrite tips at bean stage. Amphids and support cells are labeled and tinted to allow individual cells to be distinguished. Confirmed cell identities are shown for all cells visible in cross section. **(b)** Exploded diagram rendering of amphids and support cells based on 3D reconstruction. The details of amphid proto-dendrite protrusions are visible. **(c)** Rendering of same 3D reconstruction of the amphids viewed from an anterior angle highlighting the shared focus of protrusions (indicated by white arrow). **(d)** Montage of max projection of socket cell over three successive frames in the same *cnd-1* promoter labeled embryo as Fig. 5d. Soma marked with a star and protrusion with an arrow. Dynamic fluctuation is visible throughout the socket cell at this time period. **(e)** 2-fold stage dendrite tip. Colors match Fig. 5f,g. **(f)** Dendrite tips showing centrosomes absent at bean stage and present (indicated by arrows) at the comma and 1.5-fold stages.

**Supplemental Movies:**

**Movie 1:** Max projection movie of 8 hours of development of a *C. elegans* embryo with a ubiquitous histone marker (red) and a *cnd-1* promoter driven membrane marker labeling a subset of neurons (green). Temporal resolution of 1 frame/min. Same embryo as shown in Fig. 1b.

**Movie 2:** 3D rendering of embryo landmarks. Lineages with a majority fate for a given tissue type are colored to match tissue color code in Fig. 3.

**Movie 3:** Selective max projection movie of socket and sheath cell migration in C. elegans embryo with *cnd-1* driven membrane label. Same embryo as shown in Fig. 5d. A slightly looser cropping is chosen for context.

**Supplemental Data:**

Excel spreadsheets containing position and annotation information for bean, comma and 1.5-fold embryos.

**Supplemental Tables:**

**Table S1** EM Imaging Details

| **Stage** | **Estimated Time pfc** | **Slice Axial Orientation** | **Acquisition modality** | **Number of Sections** | **Lateral Resolution (nm)** | **Section Thickness (nm)** |
|---|---|---|---|---|---|---|
| Bean | 320 min | ventral-dorsal | FIB | 944 | 9.9 | 30 |
| Comma | 345 min | anterior-posterior | FIB | 1641 | 8.5 | 25 |
| 1.5-fold | 380 min | left-right | AT | 325 | 9.7 | 65 |
| 2-fold | 475 min | ventral-dorsal | AT | 310 | 19.4 | 85 |

**Table S2** EM Data Detailed Information

| | Bean stage | Comma stage | 1.5-fold stage |
|---|---|---|---|
| **Landmark Properties** | | | |
| **# Cells** | 576 | 578 | 601 |
| **Annotation Extent** | | | |
| **# Cells Confirmed Identity** | 148 | 231 | 242 |
| **# Cells Confirmed Tissue** | 198 | 177 | 95 |
| **Accuracy** | | | |
| **Single Cell Accuracy** | .78 | .73 | .71 |
| **Tissue Level Accuracy** | .95 | .91 | .79 |

**Table S3** FM Data Detailed Information

| | Bean stage | Comma stage | 1.5-fold stage |
|---|---|---|---|
| **Landmark Properties** | | | |
| **# Adjacencies Observed** | 6170 | 5838 | 5881 |
| **# Consistent Adjacencies** | 439 | 504 | 501 |
| **# Inconsistent LM** | 28 | 27 | 34 |

**Table S4** Details of FM Series Used to Time Neurite Breakouts

| Cell | EM time | FM time | Strain Source | Marker | Strain | Strain Genotype |
|---|---|---|---|---|---|---|
| AFD | 345 | 345 | [29] | *egl-13* | DCR6358 | olaEx3765 [DACR2605(egl-13p::egl-13(168bp)::SL2::PHD::unc-54UTR) at 50ng/µl + DACR2603(sdz-31p::GFP::NLS::unc-54UTR) at 25ng/µl + DACR218(unc-122p::RFP) at 25ng/µl]; ujIs113 [pie1p::mCherry::H2B::pie-1 3'UTR + nhr-2p::his-24::mCherry::let-858 3'UTR + unc-119(+)] II |
| AIY | 345 | 324 | [29] | *ttx-3* | DCR7844 | mgIs18 [ttx-3p::GFP] IV; olaEx4765 [DACR3254(ceh-24p::mCherry::unc-54UTR) at 50ng/uL + DACR218(unc-122p::RFP) at 30 ng/uL] |
| ALA | 345 | 347 | [46] | *ceh-10* | BV395 | lqIs4 [ceh-10 promoter::GFP + rol-6]; ujIs113; olaEx2383[pMM23::pLim-4_PHD_Super_Folder_GFP_unc54 UTR in pDEST-R4-R3, pUnc-122::RFP] |
| ASH/RIB | 320 | 318 | [46] | *cnd-1* | BV293 | zbIs3[cnd-1>PH::GFP] IV; ujIs113 II. |
| AVD | 345 | 339 | [46] | *zag-1* | BV450 | ujIs113 II; ex?(zag-1:GFP, pUnc-122::RFP) |
| AVG | 345 | 347 | [46] | *unc-86* | BV298 | kyIs262 [unc-86::myr GFP + odr-1::RFP] IV; ujIs113 II. |
| AWC | 320 | 324 | [29] | *ceh-37* | DCR6298 | olaEx3716 [DACR2541(ceh-37p::ceh-37(103bp)::SL2::PHD::GFP::unc-54UTR) at 100ng/ul + DACR218(unc-122p::RFP) at 50ng/ul]; ujIs113 [pie-1p::mCherry::H2B::pie-1 3'UTR + nhr-2p::his24::mCherry::let-858 3'UTR + unc-119(+)] II |
| RIH | 345 | 350 | [46] | *unc-86* | BV298 | kyIs262 [unc-86::myr GFP + odr-1::RFP] IV; ujIs113 II. |
| RIP | 345 | 330 | [46] | *unc-86* | BV298 | kyIs262 [unc-86::myr GFP + odr-1::RFP] IV; ujIs113 II. |
| RIV+ | 345 | 338 | [46] | *lim-4* | BV286 | ujIs113; olaEx2383[pMM23::pLim-4_PHD_Super_Folder_GFP_unc54 UTR in pDEST-R4-R3, pUnc-122::RFP] |
| RMDD | 345 | 313 | [46] | *zag-1* | BV450 | ujIs113 II; ex?(zag-1:GFP, pUnc-122::RFP) |
| RMED | 345 | 362 | [46] | *ceh-10* | BV395 | lqIs4 [ceh-10 promoter::GFP + rol-6]; ujIs113; olaEx2383[pMM23::pLim-4_PHD_Super_Folder_GFP_unc54 UTR in pDEST-R4-R3, pUnc-122::RFP] |
| SAAV | 345 | 324 | [46] | *lim-4* | BV286 | ujIs113; olaEx2383[pMM23::pLim-4_PHD_Super_Folder_GFP_unc54 UTR in pDEST-R4-R3, pUnc-122::RFP] |
| SIBD | 345 | 325 | [46] | *lim-4* | BV286 | ujIs113; olaEx2383[pMM23::pLim-4_PHD_Super_Folder_GFP_unc54 UTR in pDEST-R4-R3, pUnc-122::RFP] |
| SMDD+ | 320 | 297 | [46] | *lim-4* | BV286 | ujIs113; olaEx2383[pMM23::pLim-4_PHD_Super_Folder_GFP_unc54 UTR in pDEST-R4-R3, pUnc-122::RFP] |

**Table S5** Landmarks Used to Initialize Alignment

| **Cells** | **Bean stage** | **Comma stage** | **1.5-fold stage** | **Purpose** |
|---|---|---|---|---|
| Seam Cells | All | All | All | Provide A-P body axis and approximate twist along A-P. |
| Gut Cells | All | None | All | |
| Body Wall Muscles | None | None | All | |
| Distinctively positioned/shaped neurons: | RIH, RMDDL/R, AVL | PVQL/R | RIH, RMDDL/R, AVL,<br>ALA | Refine alignment of head,tail |
| Other distinctively positioned cells: | exc_cell,<br>p11,12, spike l/r | exc_cell,<br>p11,12 | exc_cell,<br>z4, z1 | |
| Distinctive Anterior Hypodermal Cells: | hyp1 (ABarappaapa)<br>hyp3(ABplaapaaaa)<br>hyp4 (ABarpapapa) | hyp4 x3<br>(ABarpapapa,<br>ABpraappaa,<br>ABplaappaa) | hyp4 x2<br>(ABpraappaa,<br>ABplaappaa) | Mark anterior |
| | hyp6 x2<br>(ABplaaaapa,<br>ABarpaapaa) | hyp6 x4<br>(ABplaaaapa,<br>ABarpaapaa,<br>ABplaaaapp,<br>ABarpapapp) | hyp6 x2<br>(ABplaappap,<br>ABpraappap) | Refine L-R axis in head |
| | | hyp7 x3<br>(ABarpaapap,<br>ABplaapppp,<br>ABpraapppp) | hyp7 x4<br>(ABpraapppa,<br>ABplaapppa,<br>ABplaapppp,<br>ABpraapppp) | Refine L-R axis in head |
| Distinctive Posterior Hypodermal Cells: | hyp7 x2<br>(Caappd, Cpappd) | | | Refine L-R axis in tail |
| | hyp8,9 | | hyp8,9 | Mark posterior |

Figure S1: Fluorescence Microscopy

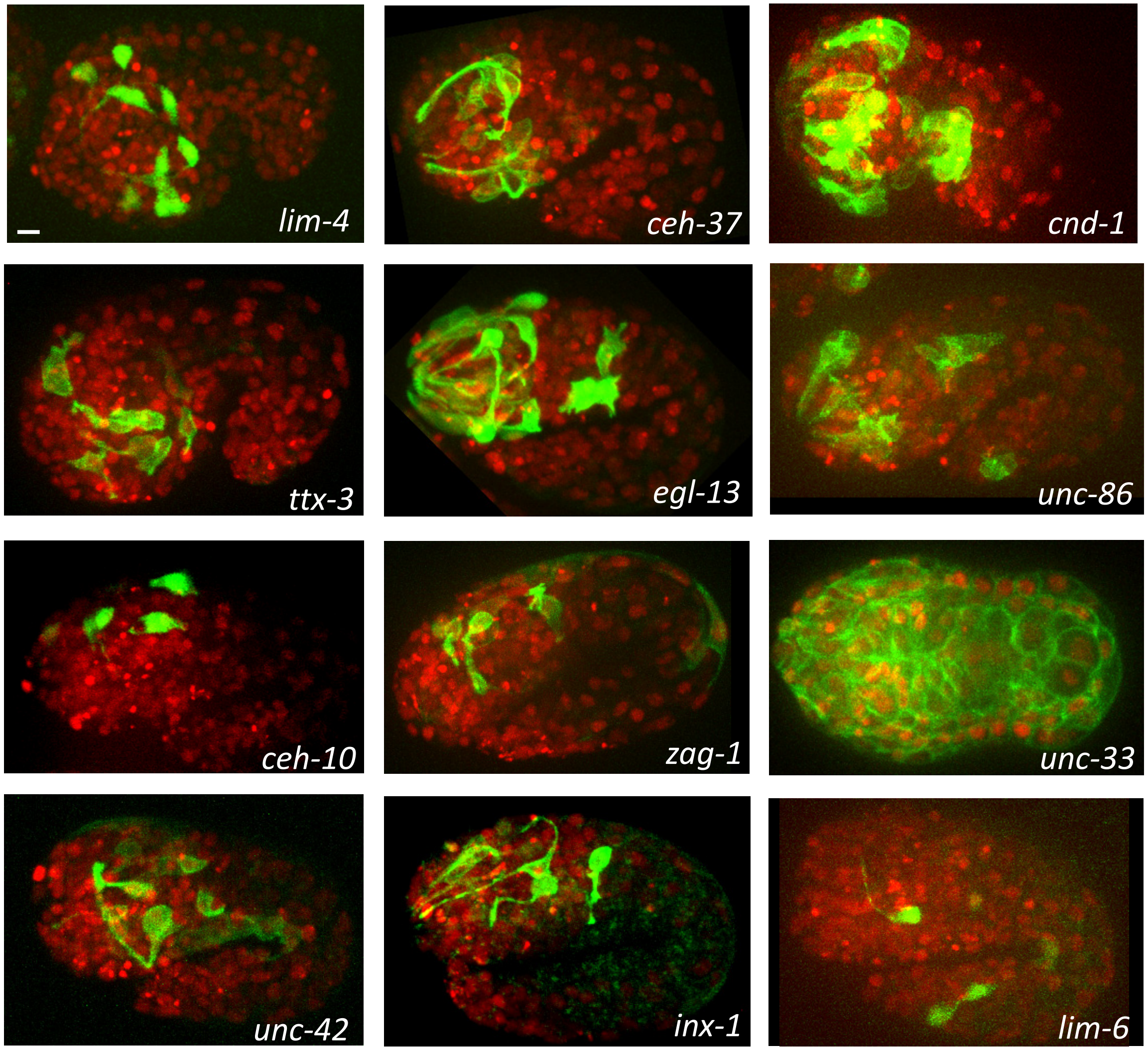

Figure 2: The Co-optimization Algorithm for Cross-modality Alignment of Developmental Data

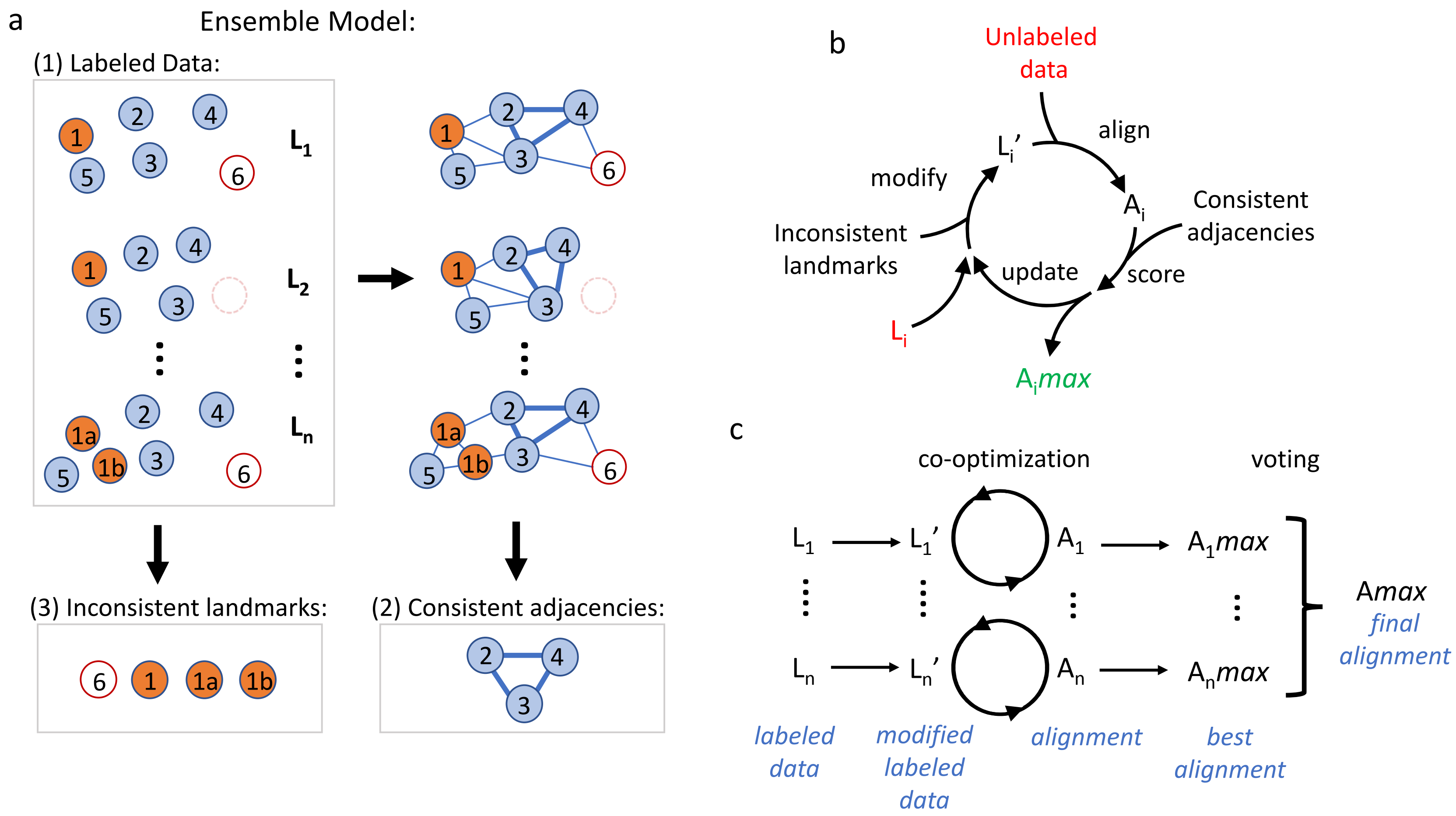

#### Figure S2: Pseudo-code of algorithm:

1. Alignment per labeled data: For each set of labeled data $L_{i=1..n}$
   - Two-step spatial pre-alignment between labeled and unlabeled data
     - Pre-Align via manually ID'd landmarks in $L_i$ and unlabeled data
     - Correspondence-less GMMCPD nonlinear pre-alignment
   - Modification and co-optimization of inconsistent landmarks:
     - $L_i$ max=$L_i$ ; $A_i$max= LAP_with_iteration($L_i$ max, unlabeled data)
     - For each inconsistent landmark present in $L_i$
       - Make modification to $L_i$ max (adding or removing landmark) creating $L_i$ max'
       - $A_i$ =LAP_with_iteration($L_i$ max',unlabeled data)
       - Score($A_i$)= missing consistent adjacencies in $A_i$
       - If Score($A_i$)< Score($A_i$max)
         - $L_i$ max = $L_i$ max'; $A_i$max = $A_i$
2. Voting: For each landmark j in unlabeled data
   - Pick most common identity for landmark j over the set of identities in $A_i$max $_{i=1...n}$
   - If 75% supermajority then the identity of landmark j is considered confident
3. Reiteration: Repeat steps 1-2 above with the modifications:
   - SubstituteTPS warping using confident identities in place of pre-alignment using manual landmarks and GMCCPD
   - Confident identities in unlabeled data remain fixed and are not recomputed

### Figure S3: Methods Details

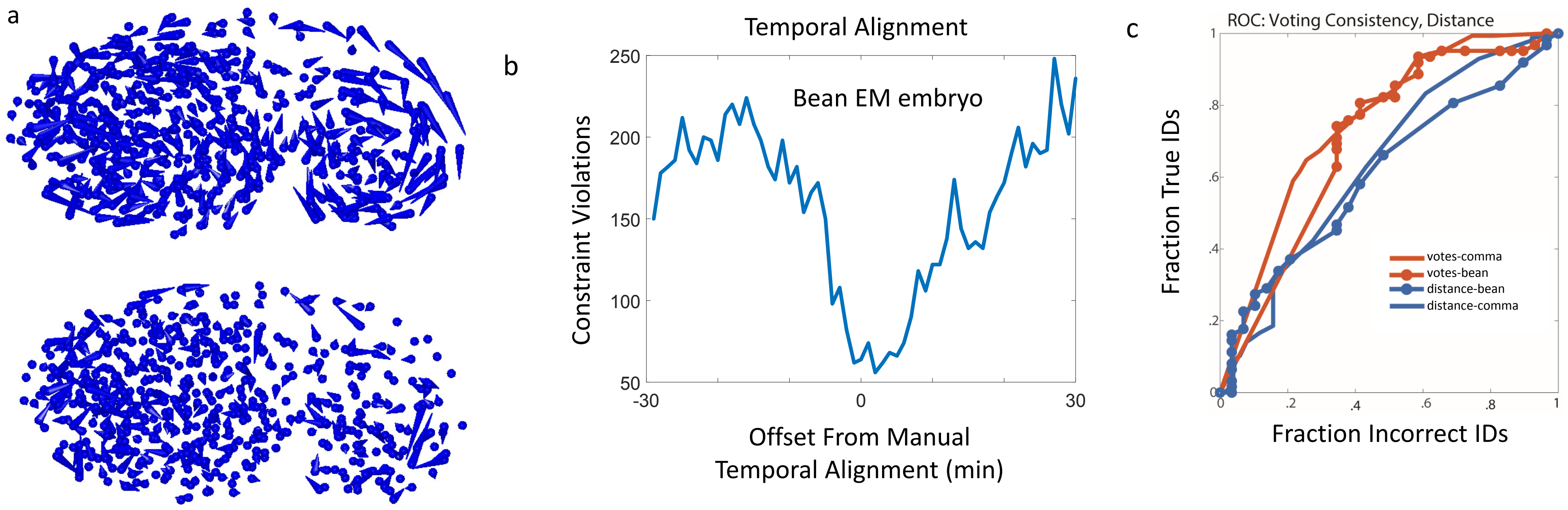

Figure S3: Cross-Modality Alignments

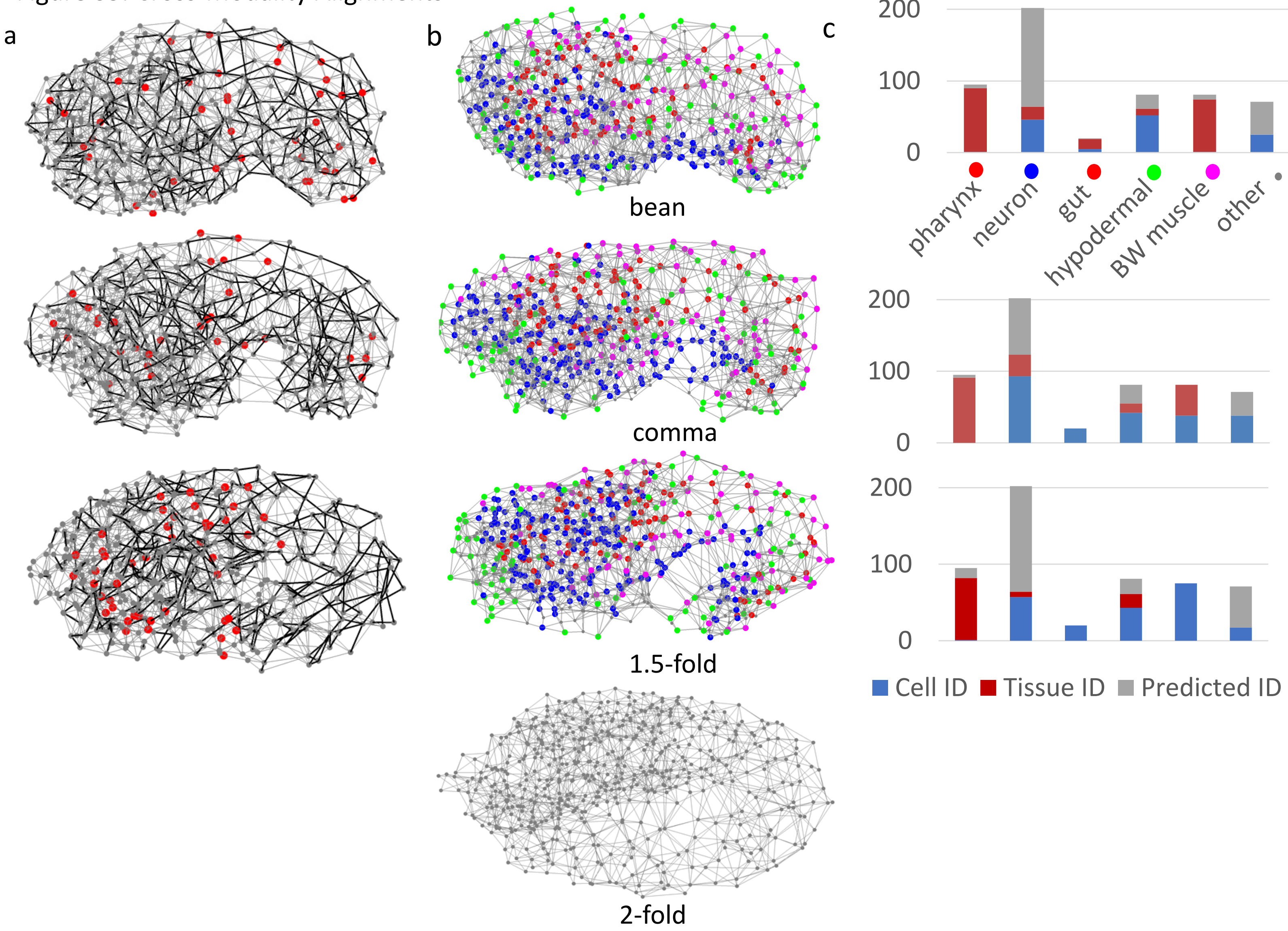

Figure S4: Earliest Neurite Outgrowths

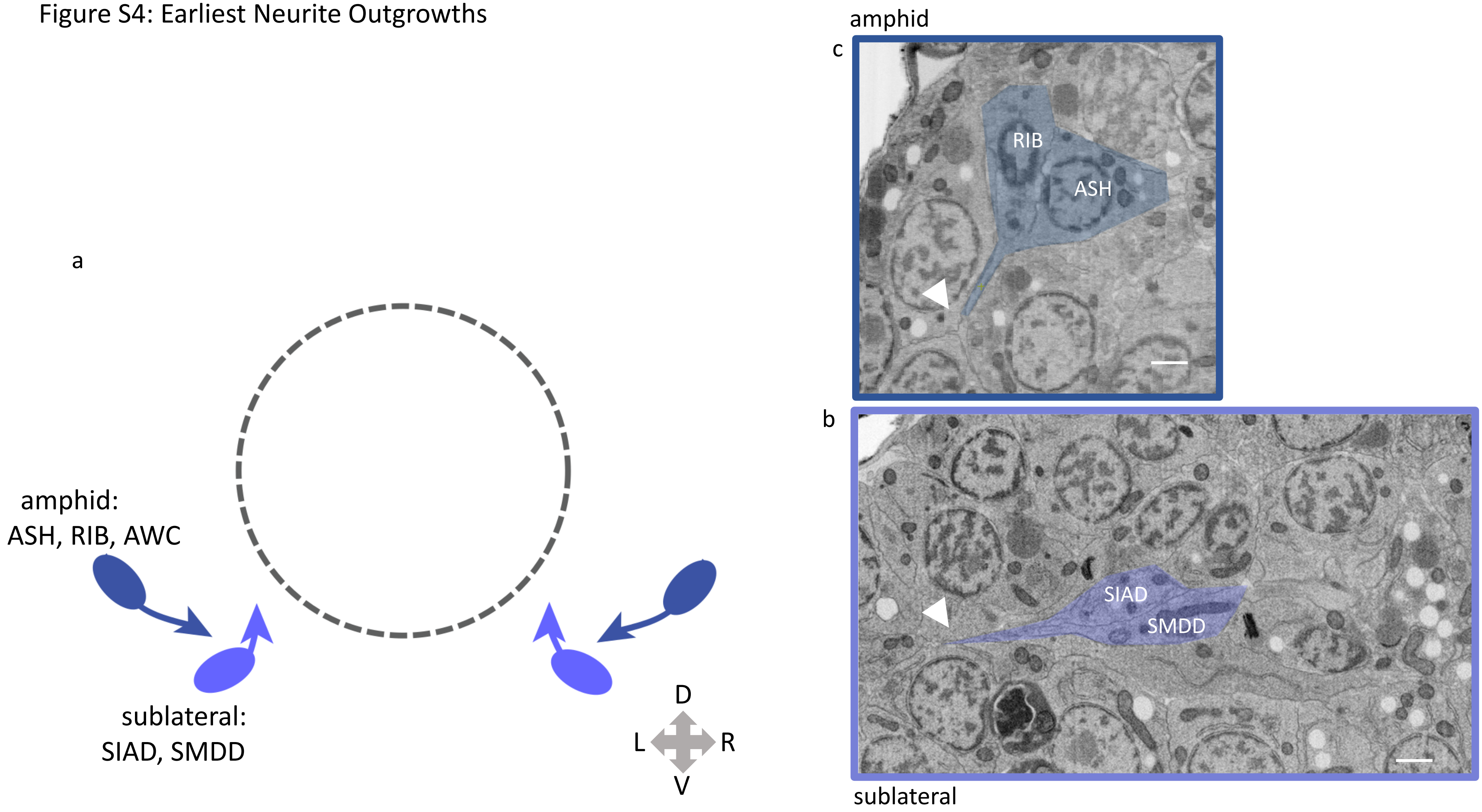

Figure S5: Pioneer and Other Early Neurites

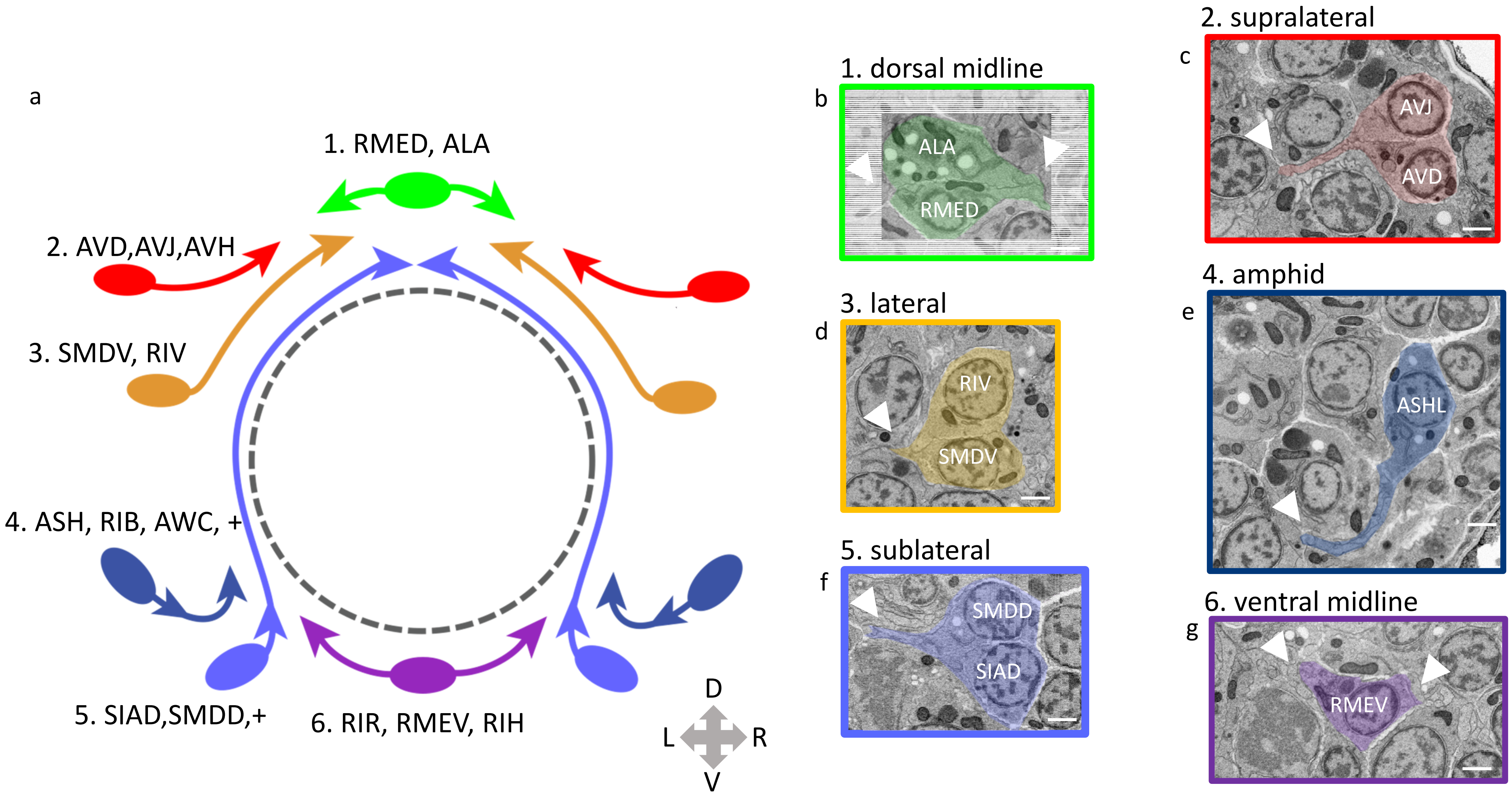

Figure S6: Amphid Organogenesis

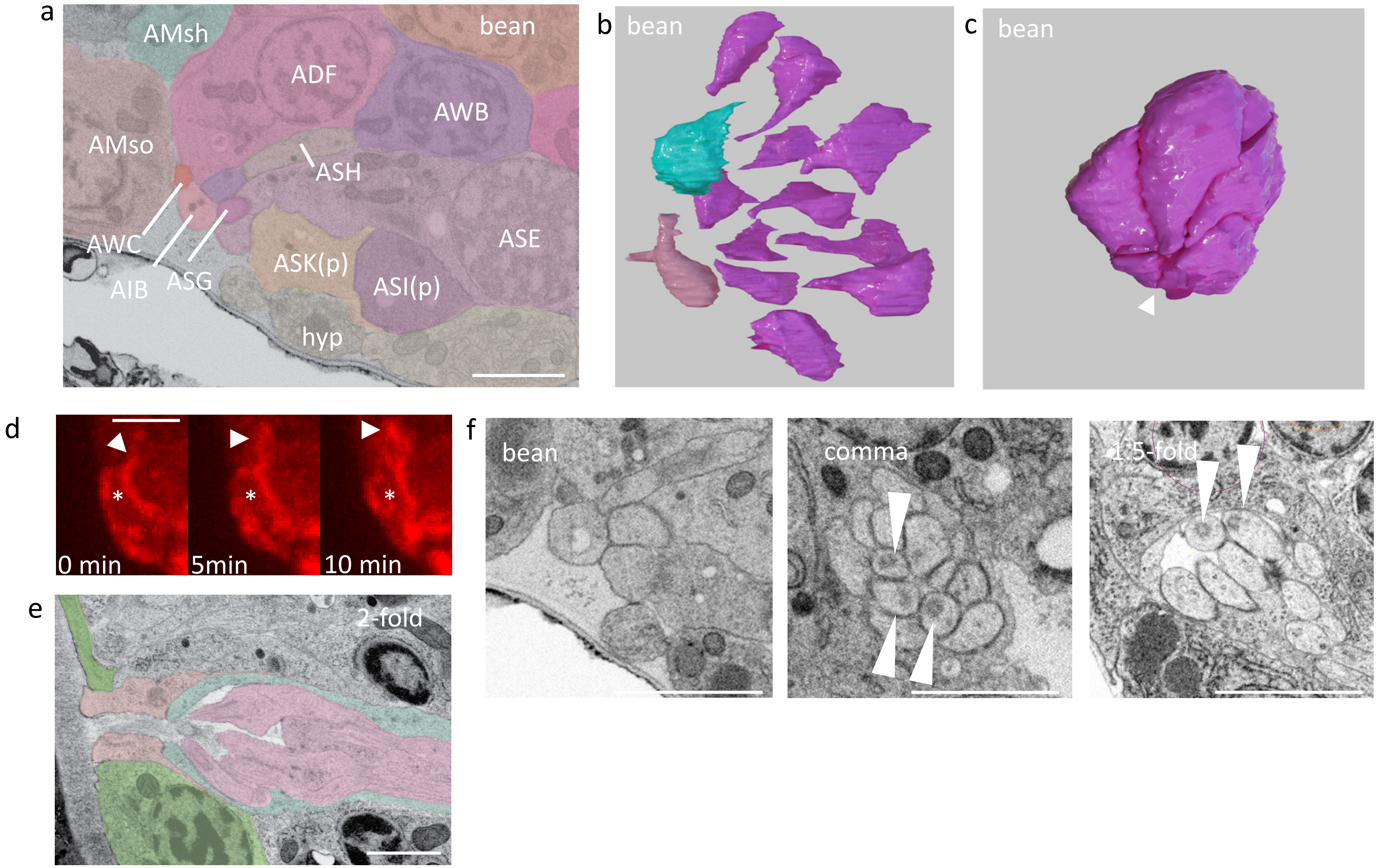
